# Multiplexed Pan Soluble Ligandome Assaying via OASIS

**DOI:** 10.64898/2025.12.21.695830

**Authors:** Yi-Hung Lee, Yesh Doctor, Yifan Zhang, Satheesh Kumar, Joseph Rainaldi, Emily Pan, Prashant Mali

## Abstract

Screening soluble protein ligands is essential for understanding signaling interactions and enabling drug discovery. Currently, screens require arrayed formats because ligand diffusibility causes non-cell-autonomous effects. To enable multiplexed pooled assaying, we developed <u>O</u>bligate <u>A</u>utocrine <u>S</u>ignaling <u>I</u>n situ <u>S</u>creening (OASIS). Using lentiviral delivery of genetically barcoded ligands fused to a tethering domain, we anchor proteins to the expressing cell’s outer membrane, exclusively enforcing autocrine signaling. Validation using IFNA2 and EGF demonstrated uncompromised autocrine signaling with significantly reduced paracrine activity. We leveraged OASIS to perform a pan-ligandome fitness screen of all 770 validated human ligands in KOLF2.1J hiPSCs, identifying potent self-renewal factors, including FGF family ligands, which we experimentally validated in soluble form. Finally, we performed single-cell Perturb-Seq to map the transcriptional remodeling induced by the pan-ligandome library. OASIS provides a generalizable framework to accelerate functional interrogation of the human ligandome and novel peptide binders within live cells and diverse lineages.

**SUMMARY:** We present <u>O</u>bligate <u>A</u>utocrine <u>S</u>ignaling <u>I</u>n situ <u>S</u>creening (OASIS), a platform enabling pooled assaying of soluble ligands by tethering genetically barcoded proteins to the surface of their expressing cells, enforcing autocrine signaling. We demonstrate OASIS by assaying all verified human ligands in hiPSCs, measuring fitness and transcriptional impacts. Our results recapitulate canonical interferon and FGF mediated signaling, validating the application of OASIS for rapid interrogation of ligands and engineered binders.

## INTRODUCTION

Receptor–ligand interactions are fundamental orchestrators of cellular homeostasis, fate specification, and pathological remodeling. Because many therapeutics—including monoclonal antibodies and cytokine mimetics—function by modulating these interactions, scalable methods to systematically map ligand activities are essential for drug discovery(*1*, *2*). Traditionally, ligand screening has relied on arrayed formats to accurately resolve individual cellular perturbations(*3*), but these approaches are cost-prohibitive and require specialized equipment.

Existing platforms have attempted to circumvent this by using CRISPRa-mediated receptor overexpression to identify ligands for orphaned receptors(*4–6*). However, these methods typically require individual screens for each ligand tested and do not easily scale to the entire ligandome. Recently, foundational work on multiplexed overexpression screens have both established methods for resolving individual library components and revealed important insights into physiology and pathology(*7*, *8*), but their application to secreted, soluble ligands remains fundamentally limited. Unlike cell-intrinsic perturbations, the diffusibility of soluble ligands creates non-cell-autonomous effects. In a pooled context, a cell is perturbed not only by its assigned ligand (autocrine signaling) but also by ligands secreted by neighboring cells (paracrine signaling), which obscures the specific linkage between a genetic perturbation and its observed phenotype.

We thus sought to utilize fusion-tethering of ligands to the cell surface to amplify autocrine effects while abrogating long-range paracrine signaling, thereby enabling high-fidelity pooled screens(*9*, *10*). Leveraging this, here, we present <u>O</u>bligate <u>A</u>utocrine <u>S</u>ignaling In situ <u>S</u>creening (OASIS), a platform for the multiplexed interrogation of soluble protein functions. We first validate that OASIS maintains potent autocrine signaling while eliminating paracrine noise. We then demonstrate its utility by performing the first pan-ligandome fitness screen in human induced pluripotent stem cells (hiPSCs), evaluating 770 human receptor-binding ligands across diverse media conditions. Finally, we integrate OASIS with single-cell Perturb-Seq to systematically map the transcriptional remodeling induced by the human ligandome at single-cell resolution.

## RESULTS

### Validation of the OASIS (Obligate Autocrine Signaling In situ Screening) Platform

We hypothesized that protein ligands fused to a C-terminal tethering domain would anchor and cluster in high density on the cell surface, facilitating efficient interaction with their cognate receptors to trigger signaling(*11*, *12*). This physical tethering would consequently restrict intracellularly produced ligands to participate exclusively in autocrine signaling, as they would be unable to diffuse away from the expressing cell membrane to influence neighbors (**Fig. 1A**). Previous studies have established that the transmembrane domain of the human CD8 alpha chain (henceforth referred to as CD8TM)(*13*) is effective for the surface display of chimeric antigen receptors (CARs)(*14*), antibody fragments(*15*), and cytokine receptors(*16*). Thus, we sought to characterize the ability of CD8TM to abrogate paracrine effects in the context of soluble ligand screening.

**Figure 1:**
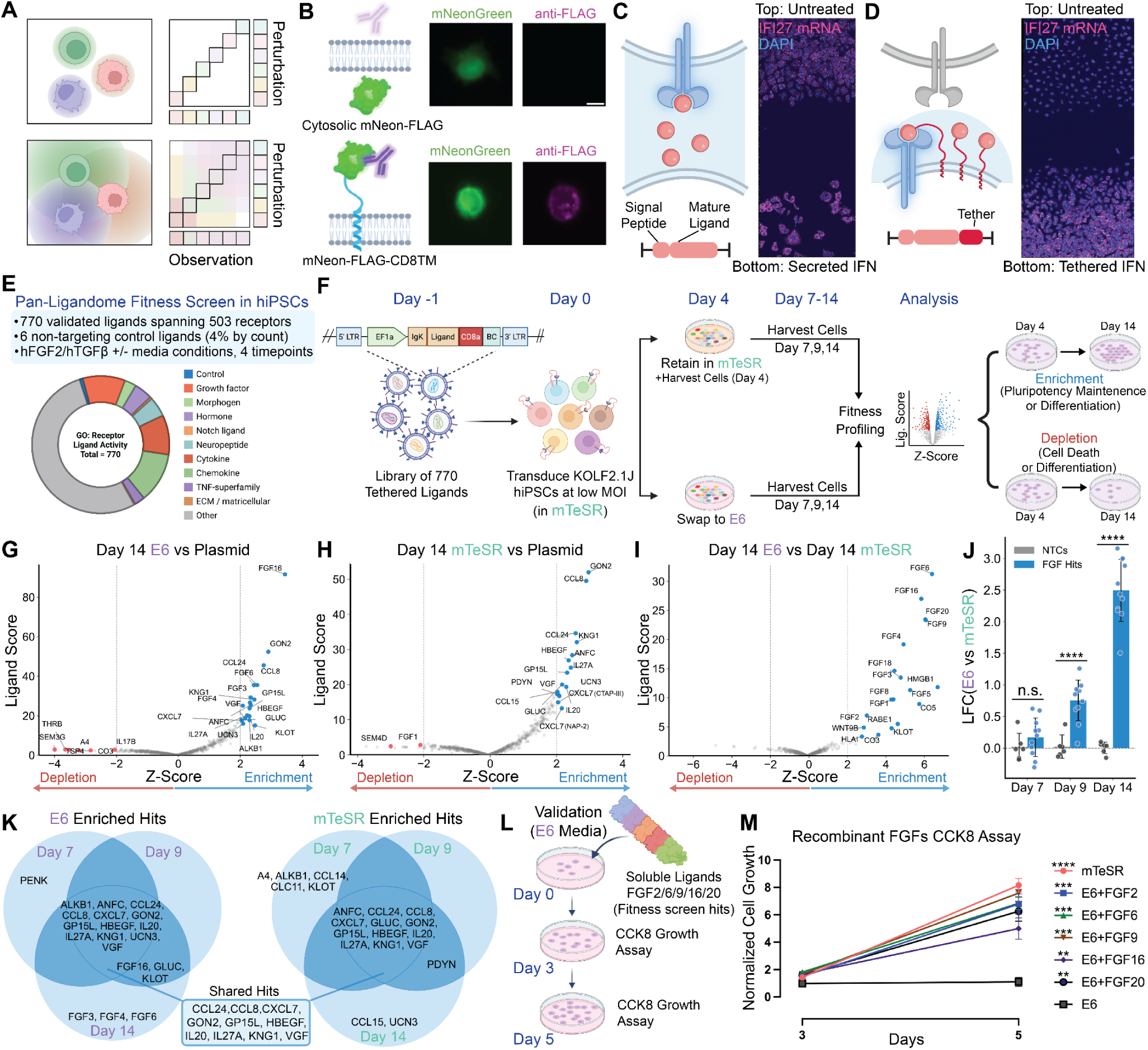
Pan-Ligandome Fitness Screen in KOLF2.1J hiPSCs via OASIS. **(A)** Schematic of autocrine signaling (top) and paracrine signaling (bottom), demonstrating how signaling perturbation is confounded and affects resolution by paracrine effects in a pooled screen context. **(B)** Immunofluorescence images of HEK293FT cell expressing untethered cytosolic mNeon (top) and CD8TM surface tethered mNeon (bottom). **(C)** Schematic of how diffusivity of secreted ligands may incur non-cell-autonomous effects (left), and experimental demonstration of this principle (right). A549 cells constitutively secreting IFNA2 (bottom) induce IFI27 response in untreated A549s (top) despite physical separation as assayed by RNA-FISH. **(D)** Schematic of tethered ligand approach (left) to abrogate non-cell-autonomous effects, and experimental demonstration of this principle (right). A549 cells constitutively secreting IFNA2 fused to a CD8TM domain (bottom) limit IFI27 response in untreated A549s (top) as assayed by RNA-FISH. **(E)** Pan-ligandome KOLF2.1J hiSPC fitness screen features. The pooled screen spans all verified Uniprot ligands, encompassing 770 unique ligands targeting 503 unique receptors, across 4 different time points and two different media conditions. **(F)** Experimental schematic. On Day 0, KOLF2.1J hiPSCs in mTeSR media were transduced with the screening library. At day 4, one plate was harvested to serve as a reference. All other plates were either maintained in mTeSR (top) or switched to E6 (bottom), and plates were harvested at days 7, 9, and 14 for both media conditions, enabling downstream fitness profiling. All conditions were performed in biological duplicates. **(G, H, I)** Volcano plot of fitness screen results using the MaGeCK test analysis. (G) Results from comparison of ligand barcode counts from day 14 in E6 vs plasmid library. (H) Results from comparison of ligand barcode counts from day 14 in mTeSR vs day plasmid library. (I) Results from comparison of ligand barcode counts from day 14 in E6 vs day 14 in mTeSR. X-axis represents the Z-score of Log Fold Changes (LFCs) across all screened ligands. Y-axis represents the score output by the MaGeCK test analysis. Blue dots indicate significantly enriched hits (Z-score > 2, FDR < 0.05) while red dots represent significantly depleted hits (Z-score < -2, FDR < 0.05). **(J)** Time course analysis of LFCs vs day 4 of enriched hits belonging to fibroblast growth factor (FGF) family and NTC ligands. (****=p<0.0001, Welch’s T-Test) **(K)** Overlap between significantly enriched ligands across E6 and mTeSR across day 7, 9, and 14 timepoints. **(L)** Schematic of validation experiment in E6 media. Recombinant (i.e. non-tethered) FGFs are added to KOLF2.1J hiPSCs on day 0 at 100ng/mL and allowed to propagate. CCK8 growth assay is performed on day 3 and day 5. Experiment performed in biological duplicate. **(M**) Results from CCK8 growth assay, normalized to E6 alone condition. (****p<0.0001, ***p<0.001, **p<0.01, two-factor ANOVA)

We first confirmed that fusion to CD8TM facilitates the surface display of ligands that are not natively trafficked to the membrane. Using mNeon-FLAG as a model (which lacks native secretion signals), we observed in HEK293FT cells that while untethered mNeon-FLAG remained intracellular, CD8TM-anchored mNeon-FLAG was successfully displayed on the outer cell membrane (**Fig. 1B**). In an orthogonal study using HEK293T cells expressing a SpyCatcher ligand tethered to CD8TM, incubation with a cell-impermeable SpyTag-FITC probe revealed clear and robust membrane localization via immunofluorescence imaging (**SI Fig. 1A**).

We next sought to answer two key questions: (1) whether C-terminal membrane tethering would compromise ligand ability to induce downstream signaling and (2) whether membrane tethering was sufficient to avoid nonspecific paracrine effects. We selected Interferon Alpha 2 (IFNA2)—a cytokine with well-defined signaling responses—as our model. In A549 cells, we conducted a transwell experiment where cells expressing either secreted IFNA2 (control) (**Fig. 1C**) or CD8TM-bound IFNA2 (**Fig. 1D**) were seeded in the bottom well, with wild-type (WT) A549s in the top well. While WT cells in the secreted control exhibited a robust interferon-stimulated response (measured by IFI27 expression), WT cells in the CD8TM-bound condition showed no such response, indicating a successful depletion of paracrine effects. Critically, autocrine signaling was preserved in the cells actually expressing the CD8TM-bound IFNA2.

Additional co-culture experiments were performed by mixing A549 cells expressing CD8TM-FLAG-bound IFNA2 with WT cells overnight (**SI Fig. 1B**). qPCR validation confirmed significant upregulation of immune markers IFI27, MDA5, and ISG15 exclusively in ligand-expressing cells (**SI Fig. 1C**). Immunofluorescence imaging further revealed that while soluble IFNA2 caused non-specific Stat2 signaling activation across the pool, the tethered IFNA2 restricted Stat2 activation to the FLAG+ ligand-expressing cells (**SI Fig. 1D**). Similar results were obtained using an Epithelial Growth Factor (EGF) ligand to induce localized EGFR phosphorylation (**SI Fig. 1E**). Collectively, these data validate that OASIS minimizes paracrine noise while maintaining potent autocrine signaling.

### Pan-Ligandome Fitness Screening in KOLF2.1J hiPSCs

Identifying soluble factors that influence human induced pluripotent stem cell (hiPSC) self-renewal and lineage commitment remains a central challenge in stem-cell biology, with direct implications for stem cell maintenance and controlled differentiation protocols. Our initial demonstration of OASIS indicated that, in principle, multiple ligands could be simultaneously interrogated without risk of nonspecific paracrine effects. We therefore sought to apply OASIS to screen for factors influencing proliferation and differentiation of hiPSCs in a pooled, multiplexed fashion. We generated a pan-ligandome fitness library **(Fig. 1E)** to screen all verified human receptor binding ligands from the Uniprot(*17*) database, totaling 770 unique full-length ligands targeting 503 unique cognate receptors. As hiPSC identity is highly influenced by media conditions and readout timepoint, we performed the screen in both media containing self-renewal factors hFGF2 and hTGFβ (mTeSR) and factor-free media (E6) conditions, across four different timepoints. To facilitate readout of ligand identity, we derived a custom vector from the mORF(*18*) backbone (see **Methods**) wherein at the protein level every ligand was tethered to the cell membrane via a CD8TM domain, and at the RNA level every ligand could be identified by a unique 20-mer RNA barcode. Collectively, this represents the first pan-ligandome pooled screen in any cell lineage.

The experimental protocol for our screen (**Fig. 1F**, see **Methods** for full details) involved initial seeding of KOLF2.1J hiPSCs at low density in mTeSR media with ROCK inhibitor (Y-27632) to ensure each cell was in a single, isolated colony to minimize downstream paracrine effects. Subsequently, cells were transduced with the pan-ligandome library at low (∼0.3) MOI and underwent puromycin selection. On day 4, plates were harvested to serve as a reference population. Non-harvested plates were then either retained in mTeSR media or transferred to E6 media, where plates thereafter were harvested at days 7, 9, and 14. All media-timepoint conditions were performed in biological duplicates. Data analysis was then performed using MAGeCK(*19*) to identify ligands that enriched cell fitness (indicating pluripotency maintenance or possible differentiation) and depleted cell fitness (indicating cell death or possible differentiation).

We confirmed high replicate correlation between biological duplicates of each condition (**SI Fig. 1F)**. We observed from the E6 (day 14) screen that cells receiving ligands from fibroblast growth factor (FGF) family members FGF3, FGF4, FGF6, and FGF16 were significantly enriched (**Fig. 1G**). These same hits were not observed in the mTeSR (day 14) conditions, an observation in line with expectations, as mTeSR already contains soluble FGF2 growth factor (**Fig. 1H**). This distinction is further underscored when directly comparing E6 (day 14) to mTeSR (day 14) conditions (**Fig. 1I**), wherein the same FGF hits are observed, with the addition of FGF1, FGF2, FGF5, FGF8, FGF9, FGF18, and FGF20. These same FGF hits are also observed when comparing E6 (day 14) to mTeSR (day 4), and absent when comparing mTeSR (day 14) to mTeSR (day 4). **(SI Fig. 1G, H)**

We further observed that FGF family induced hiPSC proliferation is time-dependent, with FGF hits having no significant effect (relative to control) at day 7, but significant effects at day 9 and 14 when comparing between the two media conditions (**Fig. 1J)**. Shared across Day 7, 9, and 14 time points across both media conditions are CCL24, CCL8, CXCL7, GON2, GP15L, HBEGF, IL20, IL27A, and VGF, indicating factors that are pro-proliferative regardless of media condition (**Fig. 1K)**. On the depleted side, semaphorin family ligands, such as SEM3C and SEM3G, have significant effects in both E6 and mTeSR conditions (**SI Fig. 1I, J**). Together, these results demonstrate the potential of OASIS for pooled ligand screening, as (1) if the screen were dominated by nonspecific paracrine effects, a strong replicate correlation between biological replicates would not be observed, and (2) if ligand tethering abrogated autocrine ligand activity, differences between FGF activity in each media condition would not be observed.

To experimentally validate our findings, we selected FGF family members identified as top hits for follow-up analysis (**Fig. 1I**). FGF2 is well established as a critical, dose-dependent regulator of human iPSC proliferation and pluripotency maintenance (as it is the growth factor natively present in mTeSR maintenance media), providing a benchmark for comparison. Our validation experiments were designed to test whether individual ligands could promote iPSC growth under E6 culture conditions and, more importantly, whether the effects identified by the OASIS screen could be recapitulated using soluble, untethered recombinant ligands. Recombinant human FGF2, FGF6, FGF9, FGF16, or FGF20 were added individually to KOLF2.1J cells cultured in E6 medium at either a high concentration (100 ng/mL). FGF2, which is known to maintain iPSC pluripotency at high concentrations, was included as a positive control, and mTeSR medium served as an additional positive control(*20*). Cell growth was assessed using the Cell Counting Kit-8 (CCK-8) assay (**Fig. 1L**). As early as day 3, cells treated with FGF2, FGF6, FGF9, FGF16, or FGF20 exhibited significantly higher viability compared to E6 alone. By day 5, this growth-promoting effect was even more pronounced, as cells cultured in E6 without added FGFs were largely non-proliferative, whereas robust proliferation was observed in the presence of each of the tested FGFs (**Fig. 1M**). Together, these validation experiments confirm that the growth-promoting effects identified in the KOLF2.1J OASIS fitness screen can be translated into a practical application using recombinant soluble FGF ligands, including FGF2, FGF6, FGF9, FGF16, and FGF20.

### Pan-Ligandome Perturb-Seq Screening in KOLF2.1J hiPSCs

While fitness screens classically quantify factors influencing proliferation or survival, an equally important and complementary dimension of ligand activity lies in their ability to modulate cell state. Particularly in hiPSCs, extracellular ligands are known to reshape transcriptional programs, bias differentiation trajectories, and alter pluripotency-associated signaling(*21*). As of yet, systematic and unbiased functional interrogation of how the human ligandome mediates transcriptional remodeling in hiPSCs has remained a major challenge. To capture these higher-dimensional responses, we combined OASIS with single-cell Perturb-seq(*22*), enabling simultaneous assignment of ligand identity and transcriptome-wide effects in a pooled format. This approach allows multiplexed interrogation of how ligands impact hiPSC cell state and gene regulatory programs at single-cell resolution, providing a transcriptional counterpart to fitness-based phenotyping.

Using OASIS, we performed Perturb-Seq in KOLF2.1J hiPSCs in both mTeSR and E6 media conditions using the same Pan-Ligandome library as in the fitness screen (**Fig. 2A)**. In the first condition, KOLF2.1J iPSCs maintained in mTeSR medium were harvested at day 4 post-transduction and processed for scRNA-seq. In the second condition, KOLF2.1J cells were maintained in mTeSR medium until day 4 post-transduction, then switched to E6 medium until Day 6, at which point cells were harvested and processed for scRNA-seq. Across both conditions, a total of ∼97,000 unique cell barcodes were detected after QC filtering for live cells harboring a single ligand.

**Figure 2:**
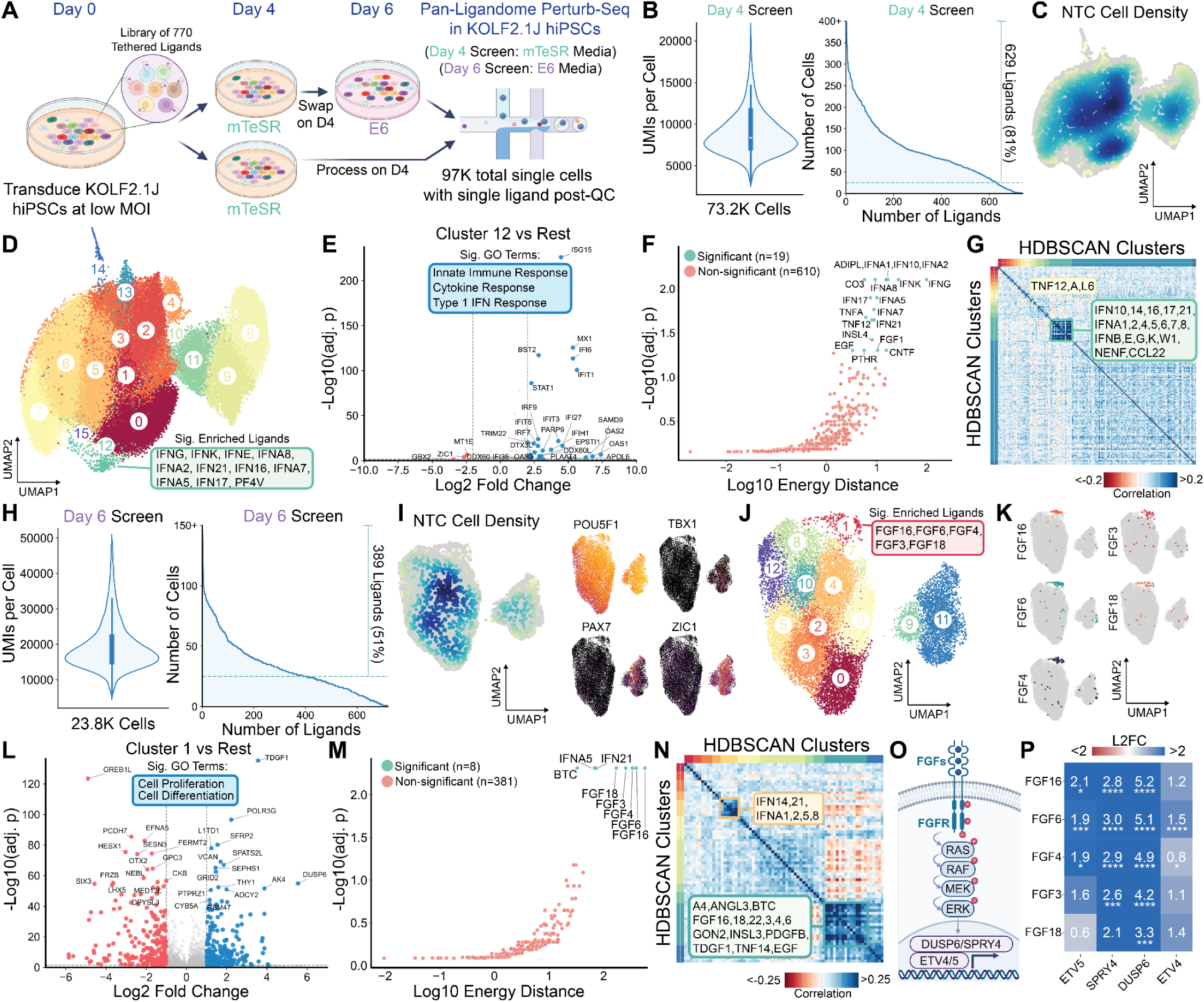
Pan-Ligandome Perturb-Seq Screen in KOLF2.1J hiPSCs via OASIS. **(A)** Schematic of Pan-ligandome Perturb-Seq in KOLF2.1J hiPSCs. Cells were transduced with our lentiviral library on Day 0 in mTeSR media. On Day 4, one plate was harvested, and single-cell RNA sequencing (scRNA-Seq) was performed, representing the Day 4 mTeSR condition. Concomitantly, a second plate was swapped from mTeSR to E6 media, allowed to propagate until Day 6, and subsequently harvested for scRNA-Seq representing the Day 6 E6 condition. Altogether, ∼97,000 single cells expressing a single ligand were identified post-QC. Panels B-G correspond to Day 4 mTeSR screen; panels H-Q correspond to Day 6 E6 screen. **(B)** Screen QC statistics for Day 4 mTeSR screen. Left: plot of UMIs per cell, representing 73.2K cells with a median of 8.5K UMIs per cell. Right: number of cells captured per ligand, with 81% of screened ligands having >25 associated cells. **(C)** Uniform Manifold Approximation and Projection (UMAP) visualization of transcriptional profiles of cells in Day 4 mTeSR screen. Gray cells represent non-NTC cells, while colored cells represent NTC cells, with the color scale demonstrating local density. **(D)** UMAP visualization colored by Leiden cluster. Cluster 12 is significantly enriched for ligands pertaining to interferon signaling, as identified by permutation testing (see Methods, adj. p < 0.05). **(E)** Volcano plot of differential gene expression between Leiden cluster 12 and all other clusters. Colored dots represent FDR < 0.05, |L2FC| > 2. Enriched GO:BP terms for upregulated genes are shown. **(F)** Energy distance test (E-test) results for all ligands vs NTC, with significant (adj. p < 0.05) ligands labeled. The E-test measures the extent of perturbation-induced transcriptional remodeling. **(G)** Clustermap of ligand-ligand transcriptional correlation for ligands identified in clusters by HDBSCAN. Insets highlight clusters related to interferon signaling and tumor necrosis factor (TNF) response. **(H)** Screen QC statistics for Day 6 E6 screen. Left: plot of UMIs per cell, representing 23.8K cells with a median of 20K UMIs per cell. Right: number of cells captured per ligand, with 51% of screened ligands having >25 associated cells. **(I)** Left: UMAP visualization of transcriptional profiles of cells in Day 6 E6 screen. Gray cells represent non-NTC cells, while colored cells represent NTC cells, with the color scale demonstrating local density. Right: marker gene expression overlays showing pluripotency (POU5F1), muscle progenitor (PAX7, TBX1), and neural (ZIC1) markers. **(J)** UMAP visualization colored by Leiden cluster. Cluster 1 is significantly enriched for ligands pertaining to FGF signaling, as identified by permutation testing (see Methods, adj. p < 0.05). **(K)** Overlay of FGF16, 4, 6, 3, 18 positions on UMAP visualization. **(L)** Volcano plot of differential gene expression between Leiden cluster 1 and all other clusters. Colored dots represent FDR < 0.05, |L2FC| > 2. Enriched GO:BP terms for upregulated genes are shown. **(M)** Energy distance test (E-test) results for all ligands vs NTC, with significant (adj. p < 0.05) ligands labeled. **(N)** Clustermap of ligand-ligand transcriptional correlation for ligands identified in clusters by HDBSCAN. Insets highlight clusters related to interferon and FGF signaling. **(O)** Mechanistic diagram of FGF-mediated signaling, indicating downstream upregulation of ETV4/5, DUSP6, and SPRY4. Modified from Ornitz and Itoh(*25*). **(P)** Log 2 fold changes (vs NTC) of downstream transcription factors in the FGF signaling pathway identified from log normalized counts from single-cell dataset. Significance is determined by differential expression via DESeq2 on unnormalized counts from single-cell dataset (see Methods, ****p<0.0001, ***p<0.001, **p<0.01, *p<0.05).

We then analyzed the OASIS Perturb-seq using the computational pipeline we had previously optimized on genome-scale perturbation data in hiPSCs(*23*) (see **Methods**) to assess the phenotypic effect of each autocrine ligand perturbation. From the day 4 mTeSR screen, we captured (post-QC) 73.2K cells with a median of 8.5K UMIs per cell, and on average 81% of screened ligands had more than 25 associated cells (**Fig. 2B**). Upon visualization of the transcriptional profiles via Uniform Manifold Approximation and Projection (UMAP), we observed pockets of the data manifold wherein NTC cells did not tend to cluster as indicated by gray regions, indicating transcriptional profiles that are potentially differentially induced by perturbing ligands (**Fig. 2C**). To quantify these effects, we first identified 15 transcriptional clusters by Leiden clustering, and found that Leiden cluster 12 exhibited significant enrichment for ligands associated with interferon signaling relative to NTCs as determined by permutation testing (**Fig. 2D, SI Fig. 2C**, see **Methods**).

Focusing on Leiden cluster 12, canonical interferon stimulated genes, such as ISG15, MX1, and BST2, were identified as the top marker genes defining this cluster (**SI Fig. 2A**). Differential gene expression analysis comparing cells in cluster 12 to all other clusters revealed robust induction of canonical interferon-stimulated genes, with genes such as ISG15 showing the most significant upregulation (**Fig. 2E**). Gene Ontology enrichment analysis of cluster 12 differentially upregulated genes highlighted pathways related to innate immune response, cytokine-mediated signaling, and type I interferon response, confirming that this cluster represents a ligand-driven interferon activation program within the hiPSC population.

When performing energy distance testing (E-test), which assesses the magnitude of perturbation-induced global transcriptional shifts, 19 ligands were identified to have significant divergence from NTCs (**Fig. 2F)**. Among those with significance were interferon ligands, including IFNG, IFNA2, IFNK, IFNL1, and IFNA1, and tumor necrosis factor (TNF) ligands TNF12, TNFA, and TNFL6. Notably, a large proportion of ligands did not induce a significant transcriptional shift, which is consistent with expectations due to competing signaling from strong pro-renewal factors hFGF2 and hTGFβ in mTeSR. We then correlated ligand-induced transcriptional profiles (**SI Fig. 2D**) and identified clusters of ligands inducing similar effects via HDBSCAN (**Fig. 2G**). Clusters of ligands related to interferon signaling and TNF response were observed, consistent with our previous findings.

Processing of the day 6 E6 screen yielded 23.8K cells with a median of approximately 20K UMIs per cell, and on average, 51% of screened ligands were represented by more than 25 associated cells (**Fig. 2H**). The Day 6 UMAP revealed two major clusters, both of which retained high expression of the pluripotency marker POU5F1. (**Fig. 2I**) Notably, the smaller cluster on the right additionally expressed markers associated with muscle progenitor identity (PAX7, TBX1) as well as neural lineage markers (ZIC1) (**Fig. 2I**). Differential gene expression analysis comparing Leiden clusters 9 and 11 (composing the cell population on the right) to all other clusters revealed enrichment of a broad and heterogeneous set of lineage-associated markers, including the neuronal marker NEUROG1 and the B-cell lineage marker PAX5 (**SI Fig. 2E**). GO term analysis of upregulated genes showed enrichment of terms associated with neural differentiation. The presence of markers from multiple and unrelated lineages indicates that these clusters may not correspond to a specific differentiation trajectory. Both clusters contained a substantial proportion of NTC cells, suggesting that the emergence of mixed pluripotency, muscle progenitor, and neural marker expression is not likely to be ligand-specific. Instead, these transcriptional states likely reflect nonspecific spontaneous differentiation induced by culturing iPSCs in E6 medium(*24*).

Leiden clustering and ligand enrichment analysis revealed that Leiden cluster 1 showed significant enrichment for ligands associated with FGF signaling, containing FGF3, FGF4, FGF6, FGF16, and FGF18 (**Fig. 2J, K, SI Fig. 2F**). Differential gene expression analysis comparing Leiden cluster 1 to all other clusters revealed enrichment of marker genes (e.g. TDGF1) and GO terms associated with cell proliferation and cell differentiation, consistent with the functional role of FGFs (**SI Fig. 2B**, **Fig. 2L**). Analysis of ligand-specific transcriptional perturbation strength, quantified by both E-test and number of DEGs per ligand, identified the same five FGFs as the top ligands inducing the largest transcriptional responses (**Fig. 2M, SI Fig. 2G**). When correlating ligand-induced transcriptional profiles (**SI Fig. 2H**) and clustering via HDBSCAN (**Fig. 2N**), clusters corresponding to interferon and FGF signaling pathways were observed.

Notably, interferon-associated transcriptional signatures were consistently detected in both the Day 4 mTeSR and Day 6 E6 scRNA-seq datasets. The presence of interferon signaling across these two distinct culture conditions indicates that the observed interferon response is a genuine ligand-driven effect rather than a secondary consequence of culture conditions or experimental variation, supporting the robustness of OASIS. We further explored whether OASIS offered sufficient resolution for mechanistic exploration or if ligand tethering abrogated downstream signaling. FGF-mediated signaling is known to induce expression of downstream transcription factors regulating proliferation and differentiation, including ETV4, ETV5, DUSP6, and SPRY4(*25*) **(Fig. 2O).** Analysis of FGF ligands enriched in Leiden cluster 1 demonstrated consistent upregulation of these canonical FGF-responsive genes. (**Fig. 2P, SI Fig. 2I-M**). The induction of these pathway-specific transcription factors further reinforces the fidelity of OASIS in capturing biologically relevant, ligand-driven signaling responses at single-cell resolution.

## DISCUSSION

In this work, we present <u>O</u>bligate <u>A</u>utocrine <u>S</u>ignaling *In situ <u>S</u>creening* (OASIS), a platform enabling multiplexed pooled assaying of soluble ligands. By combining genetically barcoded ligands with a membrane tethering domain, OASIS diminishes nonspecific paracrine effects while amplifying autocrine signaling, enabling resolution of hundreds of ligand effects in a single experiment (**Fig. 1**, top). We apply OASIS to comprehensively screen all full-length verified human receptor binding ligands in KOLF2.1J hiPSCs, identifying soluble factors significantly promoting self-renewal across 4 timepoints and 2 media conditions (**Fig. 1**, bottom). To our knowledge, this is the first pan-ligandome screen in any cell lineage. We further extend this screen to assay ligand-induced transcriptional changes via Perturb-Seq, establishing an orthogonal transcriptional readout (**Fig. 2**).

A limitation of this technology is the potential abrogation or alteration of ligand activity due to physical tethering at one end. While we demonstrated across multiple cases (e.g. IFNA2 mediated interferon response, EGF mediated EGFR activation, FGF mediated cellular proliferation) that ligands are still capable of activating their cognate receptors despite C-terminal tethering, this phenomenon may nevertheless vary on a case-to-case basis. Additionally, phenotypic alterations observed are a consequence of chronic ligand overexpression, which may not represent physiological levels of corresponding free ligands. Particularly within a screening context, downstream functional validation of candidate ligands of interest revealed from the screen in an untethered context for both specificity and activity is paramount. Finally, initial experimental optimization of cell seeding density and experimental timeline may be necessary when applying to different cell lines to ensure that separate isogenic cell colonies do not physically contact, leading to compromising paracrine signaling.

We foresee OASIS having a particular advantage in functional interrogation of peptide binders. Recent advances have been made towards reliable *in silico* candidate design of peptide binders(*26*, *27*), which require downstream *in vitro* experimental validation to assess both affinity and activity. While peptide barcoding approaches enable high-throughput testing of binding affinity, they require cognate receptor immobilization and do not offer insights into peptide activity, which requires measurement of downstream signaling consequences in live cells. OASIS presents a rapid testing platform for binders with the ability to directly assess functionality. For example, in a screen for *in silico* designed peptide cytokine mimetics, a simple flow cytometry-based readout for phospho-STAT2+ cells can be used to assess the potency of all candidate ligands simultaneously. In principle, the number of ligands that can be concomitantly screened is only limited by the number of tissue culture plates that can be maintained, enabling scaling to thousands or even hundreds of thousands of ligands. It should be noted that our pan-ligandome screen (770 ligands) was performed in a single 10cm plate per replicate condition.

In conclusion, OASIS establishes a generalizable framework for converting inherently non-cell-autonomous signaling ligands into genetically encoded autocrine perturbations. Our approach expands the searchable parameter space of receptor-ligand interactions, accelerating the discovery of endogenous ligand function, engineered binders, and factors that modulate cell state.

## ACKNOWLEDGEMENTS

We would like to thank Sami Nourreddine for useful discussions and support throughout all stages of the study. We thank Kristen Jepsen, Director of IGM Genomics Center UC San Diego. This work was generously supported by NIH grants (R01HG012351, R01NS131560), a Department of Defense Grant (W81XWH-22-1-0401), and UCSD Institutional Funds

## AUTHOR CONTRIBUTIONS

Conceptualization and Design: YHL, YD, YZ, PM; Experiments: YHL, YZ, JR, EP; Computational analyses: YD, YHL; Supervision: PM; Writing: YD, YHL, PM with input from all authors.

## METHODS

### Cell Culture

KOLF2.1Js were maintained under feeder-free conditions in mTeSR1-Plus medium (Stem Cell Technologies). Prior to passaging, tissue-culture plates were coated with growth factor-reduced Matrigel (Corning) diluted in DMEM/F-12 medium (Thermo Fisher Scientific) and incubated for 30 minutes at 37 °C, 5% CO2. For routine maintenance, KOLF2.1J were passaged 1:10 using Versene (Thermo Fisher Scientific). For lentivirus transduction and differentiation, cells were dissociated using Accutase (Stem Cell Technologies) and seeded in mTeSR-Plus with 10 μM ROCK Inhibitor Y27632 (Tocris Bioscience). All stem cells were maintained below passage 30. HEK293T cells were cultured in High-glucose Dulbecco’s Modified Eagle Medium with GlutaMax (Thermo Fisher Scientific) supplemented with 10% fetal bovine serum (Thermo Fisher Scientific A5256701) and 1x Anti-Anti (Thermo Fisher Scientific), maintained at 37°C in a humidified incubator with 5% CO₂. Cells were passed every 2-3 days upon reaching ∼80% confluence using 0.05% trypsin-EDTA (Thermo Fisher Scientific) for dissociation. HEK293FT cells were cultured in High-glucose Dulbecco’s Modified Eagle Medium with GlutaMax (Thermo Fisher Scientific) supplemented with 10% fetal bovine serum (Thermo Fisher Scientific A5256701), 0.1mM Non-Essential Amino Acids, and 1mM Sodium Pyruvate (Thermo Fisher Scientific). A549 cells were cultured in Dulbecco’s Modified Eagle Medium (Thermo Fisher Scientific) supplemented with 10% fetal bovine serum (Thermo Fisher Scientific) and maintained at 37°C in a humidified incubator with 5% CO₂. Cells were passed every 2-3 days upon reaching ∼80% confluence using 0.05% trypsin-EDTA (Thermo Fisher Scientific) for dissociation.

### OASIS Ligand Library Design and Cloning

Ligands were selected by compiling all protein entries annotated under receptor ligand activity (GO:0048018) from UniProtKB/Swiss-Prot. For each entry, the annotated peptide or chain sequence was extracted, and the corresponding signal peptide was removed prior to downstream processing. In total, 521 proteins/genes corresponding to 770 unique peptides or chains were identified, along with six control vectors expressing mCherry, SNAP-tag, Halo-tag, Cypridina luciferase, Gaussia luciferase, and β-lactamase. All amino acid sequences were codon-optimized for human expression, and each ligand peptide or chain was assigned a unique 11-bp DNA barcode. Library sequences were synthesized either as pooled oligonucleotides (<300 bp) or as gene fragments (Twist Bioscience).

To enable cell surface tethering, the CD8α transmembrane domain was incorporated as a modular anchor. Four codon variants of this domain were designed, with each variant assigned its own unique 11-bp DNA barcode. All CD8α transmembrane domain variants were synthesized as gene fragments (Twist Bioscience). The pooled ligand and anchor sequences were cloned into the TFORF-mCherry vector (Addgene #145026) under the EF1α promoter as follows: Prior to library cloning, the parental TFORF-mCherry vector was modified by removing the mCherry cassette and the TF barcode region. In their place, Igκ signal peptides flanking two BsmBI restriction sites were introduced to enable directional cloning of ligand sequences. This modified backbone is referred to as the pOASIS-library-vector.

To clone the ligand peptide pool together with their corresponding DNA barcodes, Gibson assembly reactions were prepared using 1000 ng of BsmBI-v2 (New England Biolabs)–digested pOASIS-library-vector, a 1:10 molar ratio of the oligonucleotide pool, a 1:3 molar ratio of the gene fragment synthesis pool, and a separate 1:3 molar ratio of the control gene synthesis pool. Reactions were supplemented with 2x Gibson Assembly Master Mix (New England Biolabs) and nuclease-free water to a final volume of 100 μL. Following incubation at 50 °C for 1 h, assembly products were purified using Select-a-Size MagBead (Zymo Research) with a ratio of 1.0 and electroporated into One Shot Stbl4 ElectroMAX competent Escherichia coli (Invitrogen). To assess library coverage, 1 μL of the transformation mixture was plated on LB agar plates with carbenicillin (50 μg/mL) and incubated overnight at 37 °C. The remaining transformants were expanded in 200 mL LB supplemented with carbenicillin (50 μg/mL) at 37 °C with shaking for 16 h, and the plasmid was subsequently purified using a Plasmid Maxi Kit (Qiagen). The resulting plasmid pool is referred to as pOASIS–ligand vector pool.

To clone the CD8a transmembrane domain variants as tether anchors together with their corresponding DNA barcodes, BsmBI-v2 Golden Gate assembly reactions were prepared using 800 ng of the pOASIS–ligand vector pool and 100 ng of the CD8α transmembrane domain gene fragment pool. Reactions were supplemented with BsmBI-v2, T4 DNA ligase, T4 ligase buffer (New England Biolabs), and nuclease-free H₂O to a final volume of 100 μL. Golden Gate reactions were carried out for ten cycles of 42 °C for 5 min and 16 °C for 5 min, followed by a final incubation at 60 °C for 5 min. Assembly products were purified using Select-a-Size MagBead (Zymo Research) with a ratio of 1.0 and electroporated into One Shot Stbl4 ElectroMAX competent E. coli (Invitrogen). To assess coverage, 1 μL of the transformation mixture was plated on LB agar plates with carbenicillin (50 μg/mL) and incubated overnight at 37 °C, while the remaining transformants were expanded in 200 mL LB with carbenicillin (50 μg/mL) at 37 °C with shaking for 16 h. Plasmid DNA was purified using a Plasmid Maxi Kit (Qiagen).

To assess ligand distribution, ligand-specific DNA barcodes were amplified from the plasmid pool library. The first step PCR was performed using Kapa Hifi Hotstart ReadyMix (Roche) and the primer pair OASIS-NGS-F (ACACTCTTTCCCTACACGACGCTCTTCCGATCTGTGGGCTCGGAGATGTGTAT), OASIS-NGS-R (GACTGGAGTTCAGACGTGTGCTCTTCCGATCTGCAGCGTATCCACATAGCGT). The thermocycling parameters were: 95°C for 3 min; 22 cycles of (98°C for 20 s, 65°C for 15 s, and 72°C for 20 s); and a final extension of 72°C for 5 min. Amplicons were size-selected with Select-a-Size MagBead (Zymo Research) with a ratio of 1.0x. The second step of PCR was performed using 100 ng of the first step purified PCR product.

NEBNext Multiplex Oligos for Illumina (Dual Index Primers) were used to attach Illumina adapters and indices to the samples. The thermocycling parameters were: 95°C for 3 min; 6 cycles of (98°C for 20 s, 65°C for 20 s, 72°C for 30 s); and 72°C for 2 min. Followed by purification using Select-a-Size MagBead (Zymo Research) with the ratio of 1.0, the resulting amplicon was quantified by Qubit dsDNA HS assay (Thermo Fisher Scientific) and sequenced on an Illumina NovaSeq X platform. Sequencing reads that perfectly matched each barcode were enumerated and normalized to the total number of perfectly matched reads within each condition to obtain relative ligand abundance.

### Library Lentiviral Production

HEK 293T cells were maintained in DMEM supplemented with 10% fetal bovine serum. Cells were seeded in a 15 cm plate 1 day prior to transfection, such that they were 60–70% confluent at the time of transfection. For each plate, 36 μL of Lipofectamine 2000 (Thermo Fisher Scientific 11668027) was added to 2 mL of Opti-MEM (Thermo Fisher Scientific 31985062). Separately, 3 μg of pMD2.G (Addgene #12259), 12 μg of pCMV delta R8.2 (Addgene #12263), and 9 μg of the pooled gRNA vector library were added to 2 mL of Opti-MEM. After 5 minutes of incubation at room temperature, the Lipofectamine 2000 and DNA solutions were mixed and incubated at room temperature for 20 minutes. During the incubation period, the medium in each plate was replaced with 20 ml of fresh, pre-warmed medium per well. After the incubation period, the mixture was added dropwise to each plate of HEK 293T cells. Supernatant containing the viral particles was harvested after 48 and 72 hours, filtered with 0.45 μm filters (Steriflip, Millipore). The filtered supernatant is further concentrated by adding 1 volume of lentiviral concentrator to 3 volumes of supernatant. Lentiviral concentrators are made with 40% (W/V) PEG-8000 and 1.2M NaCl, with the PH 7.0 - 7.2 and sterilized by filtering through 0.2 μM. The mixed solution was constantly rotating overnight at 4 °C. Then the solution was spun at 1600g for 60 min at 4 °C, supernatant was removed, and the viral pellet was resuspended in PBS to a final volume of 1000 μL, for each 15 cm plate. Finally, the concentrated supernatant was divided into 200 μL aliquots and frozen at −80°C. Two 15 cm plates’ worth of lentivirus was adequate.

### OASIS KOLF2.1J Fitness Screen Experiment

During the OASIS iPSC fitness screen, 3 million KOLF2.1J cells were seeded per 15-cm plate in mTeSR-Plus medium. Cells were transduced with the pooled lentiviral ligand library at a multiplicity of infection (MOI) <0.3 at day 0, followed by selection with puromycin (0.75 ng/µL) beginning two days post-transduction. At day 4, 2.5 million cells were dissociated using Accutase (STEMCELL Technologies) and replated into two 10-cm plates: one cultured in mTeSR-Plus medium supplemented with 10 µM ROCK inhibitor Y-27632, and the other cultured in E6 medium (Thermo Fisher Scientific) supplemented with 10 µM Y-27632. At least 4 million cells were harvested at this time point. Cells were subsequently passaged and harvested at days 7, 9, 12, and 14 post-transduction, with 4 million cells collected at each time point. All conditions were performed in biological duplicates.

For amplification of barcodes from genomic DNA, genomic DNA was extracted from stored cell pellets with a DNeasy Blood and Tissue Kit (Qiagen). The first step PCR was performed as two separate 100 μl reactions for each sample. 5 μg of genomic DNA was input per reaction with KAPA HiFi Hotstart ReadyMix (Roche) and the primer pair OASIS-NGS-F (ACACTCTTTCCCTACACGACGCTCTTCCGATCTGTGGGCTCGGAGATGTGTAT), OASIS-NGS-R (GACTGGAGTTCAGACGTGTGCTCTTCCGATCTGCAGCGTATCCACATAGCGT). The thermocycling parameters were: 95°C for 3 min; 22 cycles of (98°C for 20 s; 65°C for 15 s; and 72°C for 20 s); and a final extension of 72°C for 5 min. Amplicons were size-selected with Select-a-Size MagBead (Zymo Research) with a ratio of 1.0x. The second step of PCR was performed using 100 ng of the first-step purified PCR product. NEBNext Multiplex Oligos for Illumina (Dual Index Primers) were used to attach Illumina adapters and indices to the samples. The thermocycling parameters were: 95°C for 3 min; 6 cycles of (98°C for 20 s; 65°C for 20 s; 72°C for 30 s); and 72°C for 2 min. Followed by purification using Select-a-Size MagBead (Zymo Research) with the ratio of 1.0, the resulting amplicon was quantified by Qubit dsDNA HS assay (Thermo Fisher Scientific) and sequenced on an Illumina NovaSeq X platform.

### OASIS KOLF2.1J Fitness Screen Analysis

Ligand barcode amplicon libraries derived from the OASIS plasmid pool, Day 4 samples, and Day 7, Day 9, and Day 14 samples cultured in either mTeSR-Plus or E6 medium were sequenced on an Illumina NovaSeq X platform. Resulting FASTQs were analyzed using the MAGeCK Python package for computing enrichment and depletion relative to the plasmid libraries and Day 4 genomic DNA barcode abundance(*19*). Ligand scores reported in this study correspond to sgRNA-level scores output by the MAGeCK test analysis, with each DNA barcode treated as an independent sgRNA and scored individually, rather than being aggregated at the receptor level (Fig. 1H, G, I).

### OASIS KOLF2.1J Perturb-Seq Experiment

The OASIS ligand lentiviral library was delivered to KOLF2.1J cells via lentiviral transduction at low multiplicity of infection (MOI <0.3) on Day 0 in mTeSR-Plus medium. Beginning on Day 2 post-transduction, cells receiving perturbations were enriched by puromycin selection (0.75 ng/µL) through Day 6. On Day 4, a subset of cells was harvested for scRNA-seq, while the remaining cells were transitioned to E6 medium and cultured until Day 6 for additional scRNA-seq. For scRNA-seq, 100,000 transduced cells per channel from either Day 4 or Day 6 samples were loaded onto a Chromium GEM-X 3′ Chip (10x Genomics). Prior to loading, cells were dissociated using Accutase for 12 min at 37 °C and passed through a 40 µm cell strainer to obtain a single-cell suspension. Four channels were used for each time point, for a total of eight channels. Library preparation and downstream processing were performed according to the manufacturer’s instructions (10x Genomics, User Guide CG000731). Gene expression (GEX) libraries were generated using the Chromium GEM-X Single Cell 3′ Reagent Kits v4 (10x Genomics) and sequenced according to the manufacturer’s recommendations.

To map ligand barcodes to 10x single-cell barcodes, PCR amplification was performed on whole-transcriptome amplification (WTA) cDNA generated during the 10x Genomics scRNA-seq library preparation workflow. In the first PCR, 20-40 ng of WTA cDNA was used in a 100 µl PCR reaction, and a total of 25% of the WTA cDNA was used to generate ligand barcode libraries. Ligand barcodes were amplified using OASIS-Perturb-F (ACACTCTTTCCCTACACGACGCTCTTCCGATCT) and OASIS-Perturb-R (GACTGGAGTTCAGACGTGTGCTCTTCCGATCTGTGGGCTCGGAGATGTGTATAAGAGACAG).

Thermocycling conditions were 95 °C for 3 min; 14 cycles of 98 °C for 20 s, 65 °C for 15 s, and 72 °C for 30 s; followed by a final extension at 72 °C for 5 min. PCR products were size-selected using Select-a-Size MagBead (Zymo Research) at a 0.8x bead-to-sample ratio. A second PCR was performed using 100 ng of purified first-round PCR product, with NEBNext Multiplex Oligos for Illumina (Dual Index Primers) to append Illumina adapters and indices. Thermocycling conditions for the second PCR were 95 °C for 3 min; 6 cycles of 98 °C for 20 s, 65 °C for 20 s, and 72 °C for 30 s; followed by a final extension at 72 °C for 2 min. The resulting amplicon libraries were sequenced on an Illumina NovaSeq X platform.

### OASIS KOLF2.1J Perturb-Seq Analysis

Preprocessing and data analysis was performed using the same analysis pipeline as we have previously described for perturb-seq data(*23*). Briefly, Individual channels are processed using cellranger count, and aggregated per condition (Day 4 mTeSR, Day 6 E6) using cellranger aggr. Cells are assigned their corresponding barcodes using cellranger’s feature calling algorithm. Cells without exactly one assigned ligand are filtered out, with cells with >1 ligand called are likely multiplets given the low MOI of 0.3. We then perform standard single cell QC (dead cell removal, filtering out cells with low counts) using the same thresholds as we previously identified. One notable difference is that aberrant NTC cells are not filtered out via Isolation Forest filtering, as the split populations in the E6 screen resulted in all NTCs in Leiden Cluster 9/11 populations being labeled as outliers. Removing this NTC population would artificially indicate ligands in these clusters as significantly promoting cell differentiation. We then filter for ligands with at least 25 associated cells, and normalize counts per cell to 1M, followed by log1p normalization.

To analyze data, UMAP visualizations were generated using PCA on the top-2000 highly variable genes determined via the Seurat_v3 batch-aware method in scanpy(*28*) using default parameters. Embedding density plots were created using sc.tl.embedding_density with default parameters. Leiden clusters were identified using sc.tl.leiden with method = ‘igraph’ and n_iterations = 2. Cluster-level differential expression analysis was performed using sc.tl.rank_gene_groups using method=’t_test_overestimate_var’ with default parameters. Per ligand analysis including energy distance testing, differential expression testing, correlation matrix creation, and HDBSCAN clustering was performed using the same methods and parameters which we previously described(*23*).

Cluster overrepresentation analysis (Fig. 1D,K) was performed via permutation testing procedure. For each ligand, the proportion of cells in each leiden cluster was computed. Then, a random sample with replacement of the same number of NTC cells was taken, with the proportion residing in each leiden cluster being computed. This procedure was done 10,000 times. Enrichment p-values were computed as the number of times the NTC sample proportion in that leiden cluster exceeded the proportion from the ligand. All p-values were adjusted with a benjamini-hochberg procedure.

### RNA Extraction and Quantitative Real-Time PCR

RNA was extracted from cells using the RNeasy Mini Kit (QIAGEN) according to the manufacturer’s instructions. For cDNA synthesis, 1,000 ng of the extracted RNA was reverse-transcribed using the ProtoScript First Strand cDNA Synthesis Kit (New England Biolabs) following the manufacturer’s protocol. Quantitative real-time PCR (qPCR) was performed using SYBR Green Supermix (Bio-Rad) on a Bio-Rad CFX96 Real-Time PCR Detection System. Target gene expression levels were quantified and normalized relative to the housekeeping gene GAPDH, with each sample analyzed in technical duplicate, and the data were analyzed using Bio-Rad CFX Manager software.

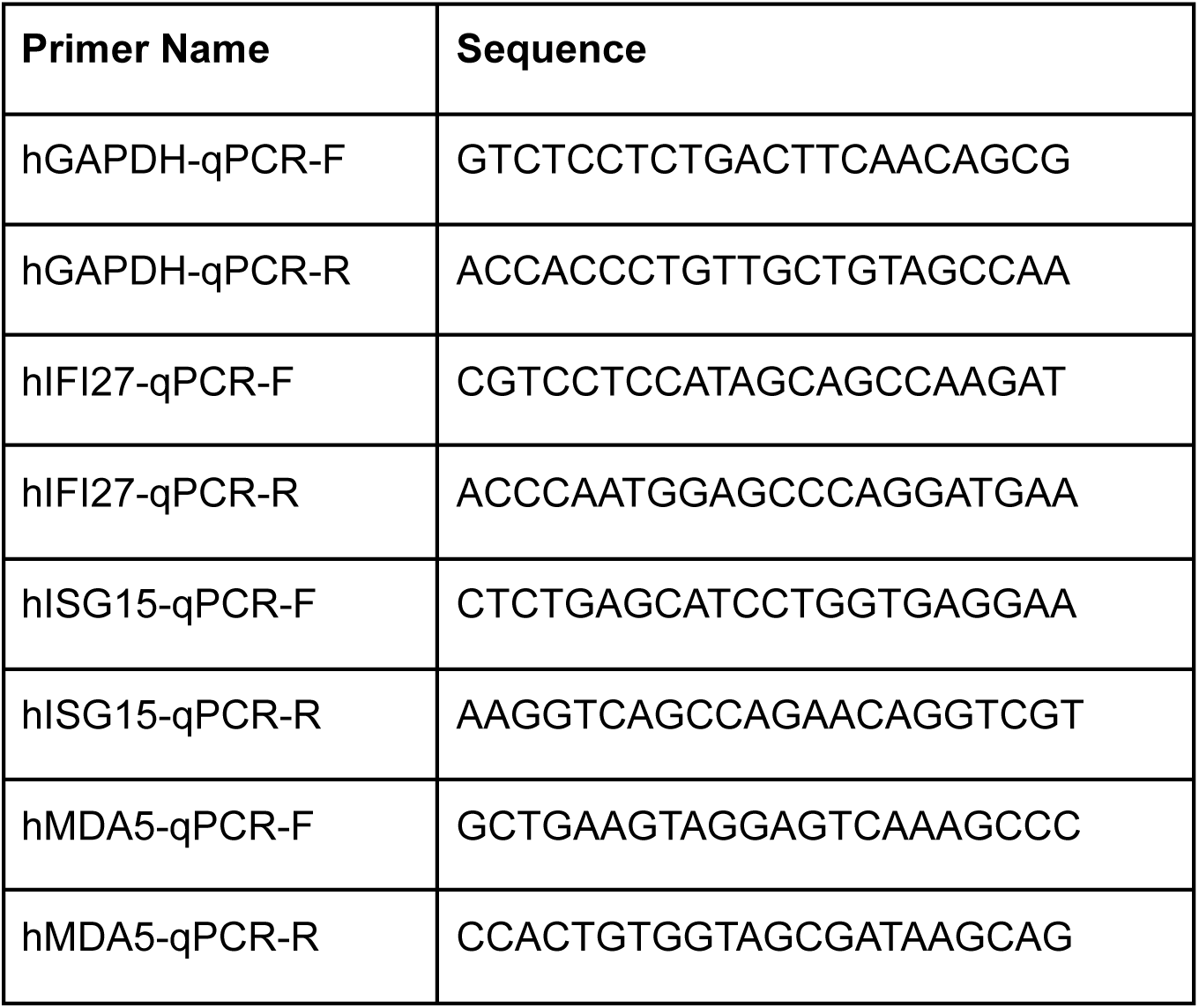

### Immunostaining and Microscopy

For immunofluorescence staining, HEK293FT or A549 cells were seeded into 96-well plates and fixed with 4% paraformaldehyde (Thermo Scientific Chemicals, 043368.9M) in PBS for 15 min at room temperature. Following fixation, cells were permeabilized with 0.1% Triton X-100 (Sigma, T8787) in PBS for 15 min, except where indicated (no permeabilization was performed for Figure 1B). Permeabilized samples were washed twice with PBS. Primary antibodies were diluted in PBS containing 4% fetal bovine serum (FBS) and applied at the following concentrations: mouse anti-FLAG (Sigma F1804, 1:500), rabbit anti–phospho-STAT2 (Cell Signaling Technology #88410, 1:100), and rabbit anti–phospho-EGFR (Cell Signaling Technology #3777, 1:400). Cells were incubated with primary antibodies overnight at 4 °C. The following day, samples were washed four times with PBS for 10 min each and incubated for 90 min at room temperature with species-appropriate secondary antibodies diluted in PBS containing 4% FBS: goat anti-rabbit Alexa Fluor 488 (Invitrogen, A11034, 1:800) and goat anti-mouse IgG Alexa Fluor Plus 647 (Invitrogen, A32728, 1:800), as applicable. Nuclei were counterstained with DAPI (Invitrogen, 62248) at 1.25 µg/mL for 10 min. Cells were then washed three additional times with PBS and stored in PBS at 4 °C prior to imaging. Raw images on the Leica DMi8 were obtained with 16 bit bit-depth per color, and highlights and shadows were adjusted in the LASX software.

### CD8-TM-SpyCatcher FITC Labeling

HEK293T cells were transduced with lentivirus encoding either SpyCatcher003-CD8 or 3x FLAG-CD8 and subsequently selected with puromycin (2 µg/mL). For surface labeling, cells were incubated with an N-terminal FITC-modified SpyTag003 peptide (FITC-MGRGVPHIVMVDAYKRYK; GenScript) diluted to 1 µM in complete growth medium for 20 min at 37 °C. Cells were gently washed twice with PBS and imaged using the Leica DMi8 and LASX software.

### Hybridization Chain Reaction (HCR™)

For the HCR™ experiment, A549 cells were transduced with lentivirus encoding either tethered or secreted form of interferon alpha subsequently selected with puromycin (2 µg/mL). Removable two-well culture inserts in 24-well plates (ibidi, 80242) were seeded with transduced cells and wild-type cells at 7 × 10^5^ cells/mL. Cells were allowed to settle overnight, after which the inserts were removed and fresh media was added. Cells were co-cultured for another 24 hours before PFA fixation and ethanol permeablization. Fixation, permeablization, and labelling were performed according to Molecular Instruments HCR™ Gold RNA-FISH User Guide. HCR™ HiFi Probes were designed and ordered from Molecular Instruments.

### CCK-8 Growth Assay

KOLF2.1J iPSCs were plated and cultured in either mTeSR-Plus medium or E6 medium supplemented with recombinant human ligands. Recombinant human FGF2 (R&D Systems, BT-FGFB), FGF6 (PeproTech, 100-30), FGF9 (PeproTech, 100-23), FGF16 (PeproTech, 100-29), or FGF20 (PeproTech, 100-41) was added to E6 medium at a final concentration of 100 ng/mL. Cell viability and proliferation were assessed on days 3 and 5 using the Cell Counting Kit-8 (Dojindo, CK04), following the manufacturer’s instructions.

---

**Supplementary Figure 1:**
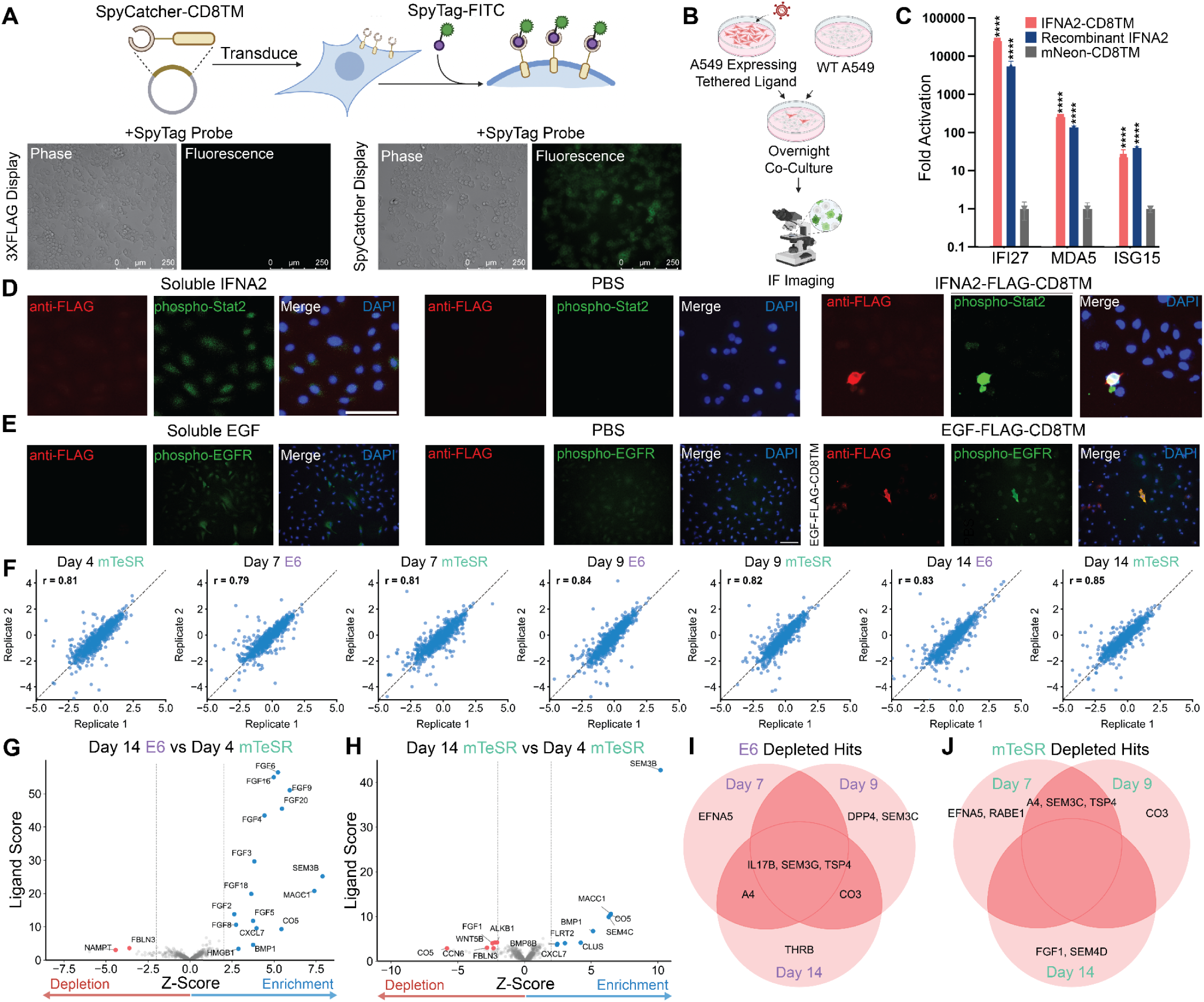
Additional Characterization of OASIS and Pan-Ligandome Fitness Screen. **(A)** (Top) Schematic of OASIS surface tethering validation in HEK293T cells. Cells transduced with SpyCatcher fused to a CD8α transmembrane domain (CD8TM) were incubated with SpyTag-FITC, which is cell-impermeable. (Bottom) Immunofluorescence (IF) images of SpyTag-FITC cell surface localization. **(B,C,D,E)** Co-culture experiment wherein A549s expressing CD8TM tethered IFNA2 (interferon-alpha 2)/EGF (epithelial growth factor) were co-cultured overnight with wild-type A549s. (B) Schematic of experimental scheme. (C) qPCR quantification of interferon signaling indicators IFI27, MDA5, and ISG15 upon activation via CD8-TM tethered IFNA2 relative to a tethered mNeon control in ligand expressing cells alone. (****=p<0.001, Welch’s T-Test). (D) Immunofluorescence (IF) images of GPI-tethered CD8TM-tethered IFNA2 with PBS/soluble IFNA2 serving as negative and positive controls respectively. anti-FLAG (red) indicates transduced cells, while phospo-STAT2 (green) indicates activated IFNAR1/2. (E) IF images of CD8TM-tethered EGF with PBS/soluble EGF serving as negative and positive controls respectively. anti-FLAG (red) indicates transduced cells, while phospo-EGFR (green) indicates activated EGFR. **(F)** Replicate correlation plots between biological replicates across timepoint and media conditions in pan-ligandome KOLF2.1J fitness screen. Axes represent LFCs relative to count in the plasmid library. **(G,H)** Volcano plot of fitness screen results using MaGeCK test analysis. X-axis represents the Z-score of Log Fold Changes (LFCs) across all screened ligands. Y-axis represents the score output by MaGeCK test analysis. Blue dots indicate significantly enriched hits (Z-score > 2, FDR < 0.05) while red dots represent significantly depleted hits (Z-score < -2, FDR < 0.05). (G) Results from comparison of ligand barcode counts from day 14 in E6 vs day 4 in mTeSR. (H) Results from comparison of ligand barcode counts from day 14 in mTeSR vs day 4 in mTeSR. **(I)** Overlap between day 7, 9 and 14 downregulated hits in E6 media condition relative to plasmid library. **(J)** Overlap between day 7, 9 and 14 downregulated hits in mTeSR media condition relative to plasmid library.

**Supplementary Figure 2:**
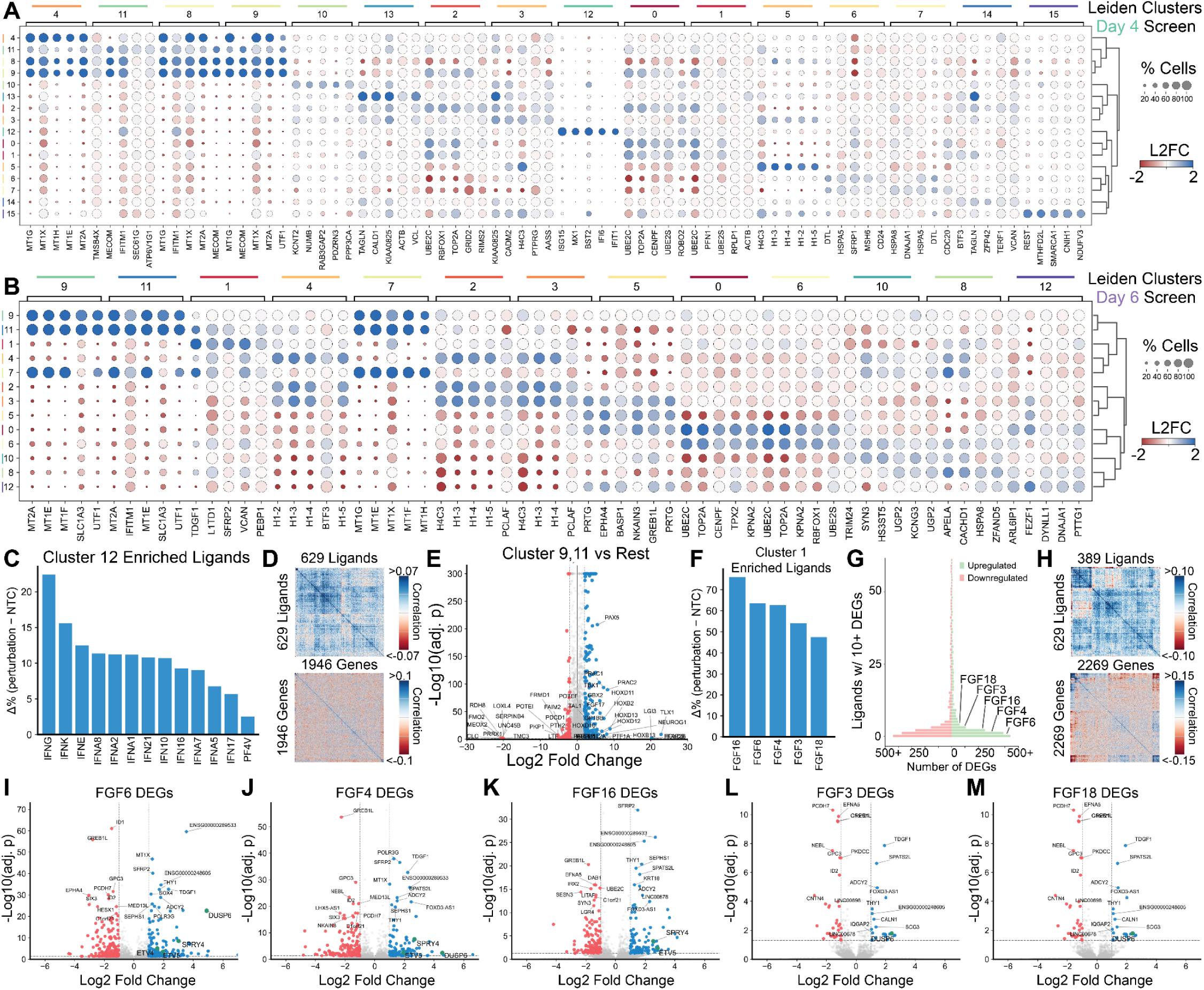
Additional characterization of Pan-Ligandome Perturb-Seq Screen. **(A)** Top-5 marker gene expression for each Leiden cluster in the day 4 mTeSR screen. **(B)** Top-5 marker gene expression for each Leiden cluster in the day 6 E6 screen. **(C)** Percent enrichment of significantly enriched ligands in Leiden cluster 12 (day 4 screen) relative to NTC. Y-axis represents the proportion delta, i.e. the % of ligand cells in cluster 12 minus the % of NTC cells in cluster 12. **(D)** Top: ligand-ligand correlation (day 4 screen) of transcriptional profiles across 1946 feature genes. Bottom: feature gene-gene correlation of expression levels across 629 ligands. **(E)** Volcano plot of differentially expressed genes in Leiden Cluster 9/11 vs all other clusters (day 6 E6 screen), with the top 20 up- and down- regulated genes by L2FC labeled. **(F)** Percent enrichment of significantly enriched ligands in Leiden cluster 1 (day 6 screen) relative to NTC. **(G)** Plot of # of differentially expressed genes per ligand, with the top 5 ligands by # of DEGs labeled. **(H)** Top: ligand-ligand correlation (day 6 screen) of transcriptional profiles across 2269 feature genes. Bottom: feature gene-gene correlation of expression levels across 389 ligands. **(I,J,K,L,M)** Volcano plot of differentially expressed genes for ligands FGF6, 4, 16, 3, and 18. ETV4/5, DUSP6, and SPRY4 are highlighted in green (if passing significance thresholds).

## REFERENCES

1. D. Yang, Q. Zhou, V. Labroska, S. Qin, S. Darbalaei, Y. Wu, E. Yuliantie, L. Xie, H. Tao, J. Cheng, Q. Liu, S. Zhao, W. Shui, Y. Jiang, M.-W. Wang, G protein-coupled receptors: structure- and function-based drug discovery. Signal Transduct. Target. Ther. 6, 7 (2021).

2. A.-H. Maehle, C.-R. Prüll, R. F. Halliwell, The emergence of the drug receptor theory. Nat. Rev. Drug Discov. 1, 637–641 (2002).

3. Parse GigaLab: 10M+ Cells in a Single Run - Redefining scRNA-seq, Parse Biosciences (2024). https://www.parsebiosciences.com/gigalab/.

4. L. Yang, T. P. Sheets, Y. Feng, G. Yu, P. Bajgain, K.-S. Hsu, D. So, S. Seaman, J. Lee, L. Lin, C. N. Evans, M. R. Guest, R. Chari, B. St Croix, Uncovering receptor-ligand interactions using a high-avidity CRISPR activation screening platform. Sci. Adv. 10, eadj2445 (2024).

5. Z.-S. Chong, S. Ohnishi, K. Yusa, G. J. Wright, Pooled extracellular receptor-ligand interaction screening using CRISPR activation. Genome Biol. 19, 205 (2018).

6. D. H. Siepe, L. T. Henneberg, S. C. Wilson, G. T. Hess, M. C. Bassik, K. Zinn, K. C. Garcia, Identification of orphan ligand-receptor relationships using a cell-based CRISPRa enrichment screening platform. Elife 11, e81398 (2022).

7. L. M. Sack, T. Davoli, M. Z. Li, Y. Li, Q. Xu, K. Naxerova, E. C. Wooten, R. J. Bernardi, T. D. Martin, T. Chen, Y. Leng, A. C. Liang, K. A. Scorsone, T. F. Westbrook, K.-K. Wong, S. J. Elledge, Profound tissue specificity in proliferation control underlies cancer drivers and aneuploidy patterns. Cell 173, 499–514.e23 (2018).

8. M. Legut, Z. Gajic, M. Guarino, Z. Daniloski, J. A. Rahman, X. Xue, C. Lu, L. Lu, E. P. Mimitou, S. Hao, T. Davoli, C. Diefenbach, P. Smibert, N. E. Sanjana, A genome-scale screen for synthetic drivers of T cell proliferation. Nature 603, 728–735 (2022).

9. J. Chen, H. S. Bernstein, M. Chen, L. Wang, M. Ishii, C. W. Turck, S. R. Coughlin, Tethered ligand library for discovery of peptide agonists. J. Biol. Chem. 270, 23398–23401 (1995).

10. C. Choi, M. N. Nitabach, Membrane-tethered ligands: tools for cell-autonomous pharmacological manipulation of biological circuits. Physiology (Bethesda) 28, 164–171 (2013).

11. A. S. Stürzebecher, J. Hu, E. S. J. Smith, S. Frahm, J. Santos-Torres, B. Kampfrath, S. Auer, G. R. Lewin, I. Ibañez-Tallon, An in vivo tethered toxin approach for the cell-autonomous inactivation of voltage-gated sodium channel currents in nociceptors: Pain modulation by tethered toxins. J. Physiol. 588, 1695–1707 (2010).

12. Y. Wu, G. Cao, B. Pavlicek, X. Luo, M. N. Nitabach, Phase coupling of a circadian neuropeptide with rest/activity rhythms detected using a membrane-tethered spider toxin. PLoS Biol. 6, e273 (2008).

13. Y. D. Muller, D. P. Nguyen, L. M. R. Ferreira, P. Ho, C. Raffin, R. V. B. Valencia, Z. Congrave-Wilson, T. L. Roth, J. Eyquem, F. Van Gool, A. Marson, L. Perez, J. A. Wells, J. A. Bluestone, Q. Tang, The CD28-transmembrane domain mediates chimeric antigen receptor heterodimerization with CD28. Front. Immunol. 12, 639818 (2021).

14. S. Rafiq, C. S. Hackett, R. J. Brentjens, Engineering strategies to overcome the current roadblocks in CAR T cell therapy. Nat. Rev. Clin. Oncol. 17, 147–167 (2020).

15. J. R. Hamilton, E. Chen, B. S. Perez, C. R. Sandoval Espinoza, M. H. Kang, M. Trinidad, W. Ngo, J. A. Doudna, In vivo human T cell engineering with enveloped delivery vehicles. Nat. Biotechnol. 42, 1684–1692 (2024).

16. C. S. Dobson, A. N. Reich, S. Gaglione, B. E. Smith, E. J. Kim, J. Dong, L. Ronsard, V. Okonkwo, D. Lingwood, M. Dougan, S. K. Dougan, M. E. Birnbaum, Antigen identification and high-throughput interaction mapping by reprogramming viral entry. Nat. Methods 19, 449–460 (2022).

17. UniProt Consortium, UniProt: The universal protein knowledgebase in 2023. Nucleic Acids Res. 51, D523–D531 (2023).

18. J. Joung, S. Ma, T. Tay, K. R. Geiger-Schuller, P. C. Kirchgatterer, V. K. Verdine, B. Guo, M. A. Arias-Garcia, W. E. Allen, A. Singh, O. Kuksenko, O. O. Abudayyeh, J. S. Gootenberg, Z. Fu, R. K. Macrae, J. D. Buenrostro, A. Regev, F. Zhang, A transcription factor atlas of directed differentiation. Cell 186, 209–229.e26 (2023).

19. W. Li, H. Xu, T. Xiao, L. Cong, M. I. Love, F. Zhang, R. A. Irizarry, J. S. Liu, M. Brown, X. S. Liu, MAGeCK enables robust identification of essential genes from genome-scale CRISPR/Cas9 knockout screens. Genome Biol. 15, 554 (2014).

20. G. Chen, D. R. Gulbranson, P. Yu, Z. Hou, J. A. Thomson, Thermal stability of fibroblast growth factor protein is a determinant factor in regulating self-renewal, differentiation, and reprogramming in human pluripotent stem cells. Stem Cells 30, 623–630 (2012).

21. P. Du, J. Wu, Hallmarks of totipotent and pluripotent stem cell states. Cell Stem Cell 31, 312–333 (2024).

22. A. Dixit, O. Parnas, B. Li, J. Chen, C. P. Fulco, L. Jerby-Arnon, N. D. Marjanovic, D. Dionne, T. Burks, R. Raychowdhury, B. Adamson, T. M. Norman, E. S. Lander, J. S. Weissman, N. Friedman, A. Regev, Perturb-seq: Dissecting molecular circuits with scalable single-cell RNA profiling of pooled genetic screens. Cell 167, 1853–1866.e17 (2016).

23. S. Nourreddine, Y. Doctor, A. Dailamy, A. Forget, Y.-H. Lee, B. Chinn, H. Khaliq, B. Polacco, M. Muralidharan, E. Pan, Y. Zhang, A. Sigaeva, J. N. Hansen, J. Gao, J. A. Parker, K. Obernier, T. Clark, J. Y. Chen, C. Metallo, E. Lundberg, T. Ideker, N. Krogan, P. Mali, A Perturbation Cell Atlas of Human Induced Pluripotent Stem Cells, Bioengineering (2024). https://www.biorxiv.org/content/10.1101/2024.11.03.621734v2.

24. Y. Lin, Embryoid body formation from human pluripotent stem cells in chemically defined E8 media. Stembook, doi: 10.3824/stembook.1.98.1 (2014).

25. D. M. Ornitz, N. Itoh, The Fibroblast Growth Factor signaling pathway. Wiley Interdiscip. Rev. Dev. Biol. 4, 215–266 (2015).

26. W. Ahern, J. Yim, D. Tischer, S. Salike, S. M. Woodbury, D. Kim, I. Kalvet, Y. Kipnis, B. Coventry, H. R. Altae-Tran, M. S. Bauer, R. Barzilay, T. S. Jaakkola, R. Krishna, D. Baker, Atom-level enzyme active site scaffolding using RFdiffusion2. Nat. Methods, 1–10 (2025).

27. M. Pacesa, L. Nickel, C. Schellhaas, J. Schmidt, E. Pyatova, L. Kissling, P. Barendse, J. Choudhury, S. Kapoor, A. Alcaraz-Serna, Y. Cho, K. H. Ghamary, L. Vinué, B. J. Yachnin, A. M. Wollacott, S. Buckley, A. H. Westphal, S. Lindhoud, S. Georgeon, C. A. Goverde, G. N. Hatzopoulos, P. Gönczy, Y. D. Muller, G. Schwank, D. C. Swarts, A. J. Vecchio, B. L. Schneider, S. Ovchinnikov, B. E. Correia, One-shot design of functional protein binders with BindCraft. Nature 646, 483–492 (2025).

28. F. A. Wolf, P. Angerer, F. J. Theis, SCANPY: large-scale single-cell gene expression data analysis. Genome Biol. 19, 15 (2018).

